# REGULATION OF HDL DYSFUNCTIONALITY BY PHOSPHATIDYLETHANOLAMINE LINKS POLY-UNSATURATED FATTY ACIDS WITH ATHEROSCLEROTIC CARDIOVASCULAR DISEASES

**DOI:** 10.1101/2025.05.27.656492

**Authors:** Malik Taradeh, Lise M. Hardy, Veronica Dahik, Marie Lhomme, Hua Wang, Canelle Reydellet, Clément Materne, KC Pukar, Eric Bun, Maud Clemessy, Jean-Paul Pais-De-Barros, Sophie Galier, Eric Frisdal, Hervé Durand, Maharajah Ponnaiah, Petra El Khoury, Elise F. Villard, Philippe Lesnik, Antonio Gallo, Laurent Kappeler, Philippe Giral, Eric Bruckert, David Masson, Maryse Guerin, Anatol Kontush, Isabelle Guillas, Wilfried Le Goff

## Abstract

**Aims:** Low plasma high-density lipoprotein (HDL)-cholesterol levels are associated with increased risk of atherosclerotic cardiovascular disease (ASCVD), potentially reflecting impaired antiatherogenic HDL functions. These latter are strongly influenced by the HDL phospholipidome, which is frequently altered in ASCVD patients. Several studies reported that plasma levels of phosphatidylethanolamine (PE) species, particularly PE (36:5), were positively associated with ASCVD, but the underlying mechanisms remain unclear. Plasma PE (36:5) exists as eicosapentaenoic (EPA)-PE and arachidonic acid (ARA)-PE, with the latter predominating in ASCVD. This study investigated whether the association of PE (36:5) with ASCVD might result from an impairment of the antiatherogenic functions of HDL.

**Methods and Results:** Total PE and PE (36:5) content of large HDL isolated from 86 women with metabolic syndrome was positively associated with carotid intima-media thickness in multivariate regression analysis adjusted for traditional risk factors. In Tg*CETP x Ldlr*^-/-^ mice fed a high-cholesterol diet, the atherosclerotic plaque size was greater when reconstituted HDL (rHDL) containing ARA-PE was injected retro-orbitally, compared with injection of control rHDL containing only phosphatidylcholine (PC). *In vitro*, PE rHDL showed reduced cholesterol efflux capacity and impaired anti-inflammatory activity in THP-1 macrophages, together with diminished anti-oxidative activity against LDL oxidation compared to control rHDL. Strikingly, ARA-PE rHDL profoundly weakened of the HDL functions, while EPA-PE counteracted the ARA-PE-induced dysfunction and potentiated the functionality of rHDL.

**Conclusion:** This study reveals a causal link between PE species, particularly ARA-PE, and HDL dysfunction, contributing to atherosclerosis. EPA-PE can restore HDL function, supporting the therapeutic potential of EPA reducing ASCVD risk.

## Introduction

Atherosclerotic cardiovascular diseases (ASCVD are the leading cause of morbidity and mortality in Western countries. It is well established that low circulating concentrations of cholesterol associated with high-density lipoproteins (HDL-C) are predictive of ASCVD incidence^1^, making HDL a promising target to reduce ASCVD^2^. However, therapeutic strategies raising HDL-C have failed to reduce ASCVD mortality^3^. Multiple biological activities are displayed by HDL, including cellular cholesterol efflux as well as anti-oxidative, anti-inflammatory, and cytoprotective activities^2^. Among them, HDL capacity to promote cholesterol efflux from macrophages is proposed to underlie the association between HDL-C and ASCVD. Indeed, the capacity of HDL to promote macrophagic cholesterol efflux is inversely associated with atherosclerosis development^4^, incident CV events^5^, and mortality in patients with myocardial infarction^6^ independently of HDL-C levels. Those studies paved the way to the notion that HDL functionality could represent a better indicator of ASCVD risk than HDL-C levels^7^.

Structure (size and shape) and composition (lipid and protein content) of HDL particles are highly heterogeneous, with such a heterogeneity being intimately linked to their biological functions^2,8^. Besides apolipoprotein (apo) A-I which constitutes around 70% of total HDL protein, proteomic and lipidomic studies identified hundreds of individual protein and lipid species in HDL^9,10^. Phospholipids account for 35-50 wt% of total HDL lipidome, with phosphatidylcholine (PC, 33–45 wt%) being the major phospholipid class^10,11^ followed by moderately abundant lysophosphatidylcholine (LPC, 0.5–5 wt%), phosphatidylethanolamine (PE, 0.5–1.5 wt%), and phosphatidylinositol (PI, 0.5–1.5 wt%)^10^. Phospholipids were reported previously to modulate HDL functionality based on their class, abundance, and biophysical properties^12–14^. Alterations of both phospholipidome and biological functions of HDL are detected in ASCVD patients, suggesting that the presence and/or the abundance of specific phospholipid species within HDL alter HDL functionality and enhance CVD risk^15–17^.

Lipidomic studies reveal positive associations between plasma levels of several PE species and ASCVD^16,18,19^. Among them, PE (36:5) displays the strongest association with ASCVD^18^, with earlier lipidomic analyses highlighting the deleterious role of this species. Thus, plasma concentrations of PE (36:5) were significantly higher in patients with stable coronary artery disease (CAD)^16^ vs controls. Even more strikingly, the prospective population-based Bruneck Study identified PE (36:5) among only three lipid species displaying the strongest predictive value for CVD^18^. Moreover, inclusion of PE (36:5) on top of traditional risk factors resulted in improved risk prediction and classification in this cohort^18^. However, molecular mechanisms through which PE (36:5) promotes ASCVD are unknown^20^.

In plasma, about 60% of total PE cargo is carried by HDL particles^11^. PE impacts structural properties of HDL^21^, and its HDL content was negatively correlated with the capacity of human serum to promote cellular cholesterol efflux^22^. Two major PE (36:5) subspecies of human plasma^23^, are PE (16:0/20:5), and PE (16:1/20:4), with the latter being speculated to predominate in cardiometabolic diseases based on the reduced eicosapentaenoic acid (EPA, 20:5) to arachidonic acid (ARA, 20:4) ratio observed in ASCVD and obesity^24,25^. EPA is an ω-3 polyunsaturated fatty acid (PUFA) that exerts several beneficial effects against ASCVD^24^. Indeed, cardiovascular risk is lower in patients receiving a highly purified EPA ethyl ester than in those on placebo^26^. A recent study indicated that circulating levels of EPA are negatively associated with inflammatory biomarkers in patients with acute myocardial infarction ^27^. Conversely, ARA, an ω-6 PUFA, is widely known for its deleterious pro-inflammatory role in ASCVD^28^. Several studies reported the capacity of PUFA to modulate HDL functionality. Hence, ω-3 fatty acids, such as EPA and α-linolenic acid, improve, while ω-6 fatty acids, such as linoleic acid (LA) attenuate the atheroprotective functions of reconstituted HDL (rHDL)^29–31^. However, effects of EPA- and ARA-containing PE (36:5) on HDL functionality remains indeterminate.

In this study, we addressed the hypothesis that PE, and particularly ARA-containing PE (36:5), can promote ASCVD by altering atheroprotective functions of HDL. We demonstrated that both total PE and PE (36:5) content in HDL was associated with atherosclerosis development in women with metabolic syndrome (MetS). We confirmed the deleterious effect of ARA containing-PE (36:5) on the ability of rHDL particles to reduce atherosclerosis in mice. Finally, our *in vitro* studies uncovered that both total PE and ARA-containing PE (36:5) attenuated the rHDL functions while EPA-containing PE (36:5) restored, or even improved them. Taken together, our study suggests that the presence of PE and ARA-PE (16:1/20:4) in HDL may represent novel biomarkers for HDL dysfunctionality, underlying the deleterious role of total PE and PE (36:5) in ASCVD. By contrast, the presence of EPA-PE (16:0/20:5) in HDL reverses this deleterious effect, providing new mechanistic clues in the understanding of the protective role of EPA in ASCVD.

## Materials and Methods

### Patients with metabolic syndrome

Carotid intima-media thickness (cIMT) was measured in a group of 86 women with metabolic syndrome (MetS) using ultrasound as previously described^32^. Mean cIMT was defined for each individual as the average of the right and the left common cIMT. Major clinical and biochemical parameters of women with MetS are presented in **Supplemental Table 1**. Patients were classified as displaying MetS on the basis of modified Adult Treatment Panel III criteria as described in our early cohort^32^. The study was performed in accordance with ethical principles outlined in the declaration of Helsinki. Written informed consent was obtained from all patients.

### Isolation and characterization of plasma HDL

HDL2 were isolated from human plasma by sequential ultracentrifugation. Briefly, the density of plasma was increased to 1.063 g/ml by addition of dry solid potassium bromide. Apolipoprotein (apo) B-containing lipoproteins (d < 1.063 g/ml) were removed after a first ultracentrifugation at 45,000 rpm for 48h at 15°C using a Beckman 50.4 rotor in a Beckman XL70 ultracentrifuge. HDL2 was isolated using a second ultracentrifugation after density adjustment of recovered bottom to 1.125 g/ml. Purified HDL2 was dialyzed using Spectrapor membrane tubing against phosphate buffer saline (PBS) at pH 7.4 before analysis for their lipid and protein content using Indiko™ Plus clinical chemistry analyzer (Thermo Scientific, US) according to the manufacturer’s instructions^12^, stored at +4°C and used in functional assays.

### Preparation of reconstituted HDL

ApoA-I was isolated from human plasma as described elsewhere (32). The purity of individual ApoA-I-containing fractions detected at 280 nm was assessed on 20% denaturing acrylamide gel revealed with Coomassie blue. Pure fractions were pooled and dialyzed against ammonium buffer (20 mM, pH 7.4) and lyophilized^33^.

#### Preparation of rHDL

rHDL particles were prepared by sodium cholate dialysis as previously described^33^ using L-α phosphatidylcholine (Soy-PC) without or with either L-α phosphatidylethanolamine (Soy-PE; Avanti Polar Lipids, AL, USA), PE (16:1/20:4), or PE (16:0/20:5; synthesized by ICBMS, Lyon, France), at a molar ratio of 1:90 (ApoA-I: Soy-PC) for Soy-PC rHDL and 1:70:20 (ApoA-I: Soy-PC:PE) for either Soy-PE, PE (16:1/20:4), or PE (16:0/20:5) rHDLs micelles (**Supplemental Table 2**). rHDL concentrations were based on their ApoA-I content, measured using Indiko™ Plus clinical chemistry analyzer (Thermo Scientific, US) according to the manufacturer’s instructions. Quality of rHDL was controlled using non-denaturing TBE 4– 20% polyacrylamide gel stained with Coomassie Brilliant Blue (**Supplemental Figure 1**)^29^.

### Lipidomic analysis

LC/MS-MS was used to characterize the phospholipidome of rHDL particles as previously described^34^.

### Atherosclerosis development and injection of reconstituted HDL in mice

Mice were housed in a conventional animal facility and fed ad libitum a normal chow diet. For the study of atherosclerosis, female Tg*CETP* x *Ldlr*^-/-^ mice (9-10 weeks of age, weight of ±22g) were fed a high-cholesterol diet (HCD) (1.25% cholesterol, 16% cocoa butter, SAFE diet N°CD002510, France) for 8 weeks. Then, mice under isoflurane anesthesia (2% isoflurane/0.2 L O_2_/min) were retro-orbitally injected with either ARA-PE rHDL (6 mice) or control Soy-PC rHDL (5 mice) at a dose of 15 mg ApoA-I/kg of body weight three times per week for a period of two weeks upon a normal chow diet.

Injection of rHDL was validated through the quantification of plasma levels of human ApoA-I after a single injection of Soy-PC rHDL (15 mg ApoA-I/kg of body weight) or buffered saline in female Tg*CETP* x *Ldlr*^-/-^ mice (**Supplemental Figure 2A**), while the HCD was validated through the quantification of plasma cholesterol levels after 8 weeks of HCD (**Supplemental Figure 2B**). The effectiveness of control Soy-PC rHDL toward atherosclerosis regression was validated in an independent study by injecting female Tg*CETP* x *Ldlr*^-/-^mice with either 15 mg ApoA-I/kg of Soy-PC rHDL or buffered saline three times per week for a period of 4 weeks upon a normal chow diet (**Supplemental Figure 2C and 2D**). Quantification of plasma protein and lipid levels was performed as previously described^12^. Mice were sacrificed by cervical dislocation and tissues were collected, snap-frozen and stored at −80°C or fixed in 10% formalin for further analysis. All procedures were approved and accredited (No. 02458.02) by the French Ministry of Agriculture and were in accordance with the guidelines of the Charles Darwin Ethics Committee on animal experimentation.

To quantify atherosclerotic plaques, hearts perfused with sterile PBS were collected and fixed in 10% formalin solution as previously described^35^ and incubated in 20% sucrose-PBS solution for 24 hours. Hearts were dissected at the level of the aortic root, embedded in Tissue-Tek optimum cutting temperature (OCT) medium (Sakura Finetek Europe, The Netherlands) and snap frozen in liquid nitrogen. 60 cryosections (10 µm thickness) were cut through the proximal aorta using Leica CM1900 Cryostat (Leica Biosystems), fixed in 10% formalin for 5 minutes and stained with filtered Oil red-O for 10 minutes. Atherosclerotic lipid lesions in the aortic root were quantified using ImageJ software (National Institutes of Health) on the images of a Zeiss AxioImager M2 microscope and plaque area measured with the AxioVision Zeiss software.

### Human macrophages

*Human macrophages.* Human THP-1 monocytes obtained from American Type Culture Collection were maintained at 37°C in 5% CO2 in RPMI 1640 media containing 10% heat-inactivated fetal bovine serum (FBS), 2 mmol/L glutamine, and 100 U/ml penicillin/streptomycin. Cells were differentiated into macrophage-like cells with 50 ng/mL phorbol 12-myristate 13-acetate (PMA) for 48-72h.

### Cholesterol efflux capacity of reconstituted HDL

The capacity of reconstituted HDL to promote cholesterol efflux from human THP-1 macrophages was evaluated as previously described^36^. Cellular cholesterol efflux to Soy-PC, Soy-PE, ARA-PE, and EPA-PE rHDL (at 5, 10, 20, and 50 μg ApoA-I/ml) was assayed in a serum-free medium for a 4-hour chase period.

### Antioxidative activity of rHDL

Antioxidative activity of rHDL (final concentration of 50µg ApoA-I/ml) was evaluated towards reference LDL (final concentration of 0.2 mg cholesterol/ml) isolated from a pool of plasma obtained from healthy subjects by the Etablissement Français du Sang (EFS). LDL was isolated from normolipidemic plasma by isopycnic density gradient as described earlier^17^. The chemical composition of LDL was determined using commercially available assays (Diasys, Germany)^17^. rHDL particles were added to LDL directly before inducing oxidation by copper sulfate (CuSO_4_, final concentration 0.05 µM). LDL oxidation was assessed using 2ʹ,7ʹ-dichlorofluorescein diacetate (DCFH) fluorescent probe by Fluorescence Microplate Reader (Gemini, Molecular Devices, USA) as previously described^37^.

### Anti-inflammatory activity of rHDL in THP-1 macrophages

Human THP-1 macrophages plated at a density of 1.0 × 10^6^ cells/well into 24-well plates were treated with different rHDLs (20 µg ApoA-I/ml) in a serum-free RPMI media for a 4-hour (early) or a 16-hour (late) period. rHDL-containing media were removed and cells were washed twice with PBS before inducting inflammation with lipopolysaccharide (LPS; 100 ng/ml) for 4 hours when indicated. After the LPS treatment, the culture media were collected and secreted Interleukin-1 beta (IL-1β) were measured by a MILLIPLEX Magnetic Bead Panel (Millipore) and the MAGPIX device (Luminex) system according to the manufacturer’s instructions.

### RNA extraction, retro-transcription and real-time quantitative PCR

Cell’s total RNAs were extracted using the NucleoSpin RNA II kit (Macherey-Nagel). Reverse transcription and real-time qPCR were performed using a LightCycler LC480 (Roche) as previously described^36^. Human mRNA levels were normalized to the mean expression of three housekeeping genes: human non-POU domain-containing octamer-binding housekeeping gene (NONO), human α-tubulin (TUBA) and human heat shock protein 90kDa alpha (cytosolic), class B member 1 (HSP90AB1). Data were expressed as a fold change in mRNA expression relative to control values. The primers used in this study are in **Supplemental Table 3.**

### Phospholipid transfer from LDL to rHDL

The transfer of phospholipids from LDL to rHDL was evaluated using human LDL labeled with Dil (1, 1’-dioctadecyl-3, 3, 3’, 3’-tetramethylindocarbocyanine perchlorate) fluorescent probe as described previously for the lipolytic assay^38^.

### Lipid surface and core fluidity of rHDL

The fluidity of the lipid surface and core of rHDLs was evaluated using trimethylamine-diphenylhexatriene (TMA-DPH), and diphenylhexatriene (DPH; Cayman Chemical, USA) fluorescent probes, respectively^39^. Reconstituted HDL particles (3.3 mg ApoA-I/dL) were incubated with DPH (final concentration of 0.80 µM, dissolved in tetrahydrofuran) or TMA-DPH (final concentration of 0.32 µM, dissolved in dimethylformamide) at 37C° for 1 hour to achieve the incorporation of the fluorescent probes into the lipoproteins. Anisotropy of the probe fluorescence was measured by FlexStation 3 Multi-Mode Microplate Reader (Molecular Devices, USA) using a standard polarizer set as previously described^39^.

### Cell surface expression of TLR4

Cell surface expression of TLR-4 was investigated using R-Phycoerythrin (PE) fluorescent labeled-mouse anti-human TLR-4 (CD284) antibody (Clone TF901, 564215; BD Biosciences, US). THP-1 macrophages treated 4h with 20 µg ApoA-I/ml of rHDL, detached with EDTA-trypsin and stained 30 minutes with 100 µl of eFluor™ 520 Fixable Viability dye (Invitrogen, US) diluted in PBS-FBS 5% (1:1000 v/v). After centrifugation (5 minutes 400 g), pelleted cells were stained 30 minutes with 100 µl of PE fluorescent-labeled mouse anti-human TLR-4 (CD284) antibody diluted in PBS-FBS 5% (1:50 v/v) with FcR blocking reagent (Miltenyi Biotec). Cells were fixed using FOXP3 fixation diluent (Invitrogen, Thermo Fisher Scientific, US) and analyzed by flow cytometry (LSR II FORTESSA SORP (BD Biosciences)).

### Western blotting analysis

Total protein from THP-1 macrophages plated at a density of 4.0 × 10^6^ cells/well into 6-well plates treated 4h with different rHDLs (20 µg ApoA-I/ml) and stimulated 30 minutes with LPS, was extracted and analyzed by Western blotting as previously described^14^ using twenty micrograms of protein. Quantification of Western blots was performed using Li-Cor scanner (Odyssey system, Li-COR Biosciences, Germany). Antibodies were purchased from Cell Signaling Technology (Massachusetts, USA) **Supplemental Table 3**.

### Eicosanoid production by LC/MS-MS

The impact of rHDLs on the long-term eicosanoids production was investigated in THP-1 macrophages plated at a density of 4.0 × 10^6^ cells/well into 6-well plates and treated with different rHDLs (20 µg ApoA-I/ml) for a period of 16h. At the end of the treatment, cells were detached with EDTA-trypsin and centrifuged, supernatant was aspirated and cell pellets were stored at −80C for LC/MS-MS analysis. Detailed procedure is in supplemental methods.

### Statistical analyses

Data are presented as mean ± S.E.M. Experiments were performed at least in triplicate and values shown represent at least three independent experiments. The normal distribution of the data was evaluated using Shapiro-Wilk and Kolmogorov-Smirnov tests. For normally distributed data, comparisons were performed by an unpaired two-tailed Student’s t-test with Welch’s correction, multiple Student’s t-test and Welch-ANOVA test without assuming identical standard deviation (SD) for the both populations, unless indicated in the graph legends. For not normally distributed data, Mann-Whitney test and Dunn-ANOVA test were used. Statistical analyses were performed by the Prism software from GraphPad (San Diego, CA, USA). The association between the abundance of PE (36:5) in large HDL2 particles and cIMT was evaluated by univariate and multivariate linear regression using the R statistical software-version 3.3.2 (R Foundation for Statistical Computing).

## Results

### Total PE and PE (36:5) contents of large HDL were associated with atherosclerosis in patients with MetS

Earlier lipidomic studies have reported that plasma levels of PE species, especially PE (36:5), are strongly and positively associated with ASCVD^16,18^. To test if these effects result from the presence of PE and PE (36:5) in HDL, the association of their contents in large particles, the primary contributors to PE cargo among plasma HDL subparticles^40^ and the main cholesterol efflux effector in women^41^, with carotid intima-media thickness (cIMT) was investigated in a cohort of 86 women with MetS. Unadjusted linear regression analysis showed that both total PE and PE (36:5) contents of large HDL2 were positively associated with cIMT (β=0.285, p=0.0087, **Figure 1A**; β=0.359, p=0.0008, **Figure 1B**, respectively). This association remained significant with total PE (β=0.308, p=0.0053, **Figure 1A**) and PE (36:5) (β=0.268, p=0.0078, **Figure 1B**) after adjustment for ASCVD risk factors including age, diabetes mellitus, hypertension, smoking, plasma total cholesterol and triglycerides levels. This result indicates that total PE and PE (36:5) content of large HDL was intimately associated with atherosclerosis in women with MetS, suggesting a potentially deleterious role of PE in the atheroprotective function of HDL.

**Figure 1.**
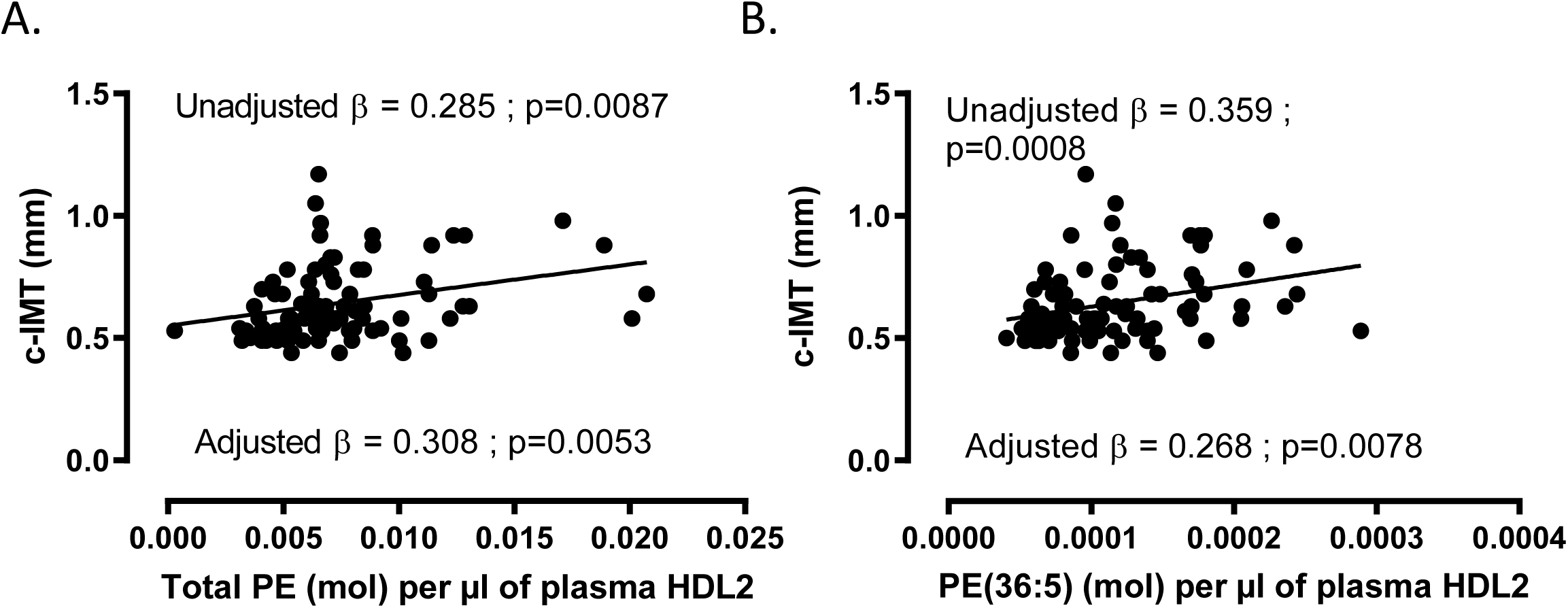
Total PE and PE (36:5) content of HDL2 was associated with atherosclerosis in women with Metabolic Syndrome. Human HDL2 particles were isolated from plasma of 86 women with Metabolic Syndrome by sequential ultracentrifugation for 48 hours. Total PE and PE (36:5) contents in HDL2 was quantified by LC-ESI/MS/MS. Carotid intima-media thickness (cIMT) was evaluated by ultrasonography and correlated with total PE and PE (36:5) content in HDL2. A. Unadjusted and adjusted linear regression analysis for the mean cIMT (in mm) according to total PE content of HDL2 B. Unadjusted and adjusted linear regression analysis for the mean cIMT (mm) according to PE (36:5) content of HDL2. The analysis was adjusted for traditional risk factors, including age, diabetes mellitus, hypertension, smoking, plasma total cholesterol and triglycerides levels. HDL: high density lipoprotein; PE: Phosphatidylethanolamine.

### Lipidomic characterization and fatty acid profiling of reconstituted HDL

To investigate the impact of PE in general, and specifically PE (36:5), on HDL functionality, we generated four types of rHDL; including Soy-PC rHDL, containing only Soy-PC as a control, and three experimental rHDLs, Soy-PE rHDL, ARA-PE rHDL, and EPA-PE rHDL, each comprising Soy-PC mixed with Soy-PE, ARA-PE, or EPA-PE, respectively, at the same 7:2 molar ratio. Soy-PC and Soy-PE were used as natural mixtures of several PC and PE species, while ARA-PE (16:1/20:4) and EPA-PE (16:0/20:5) were used as the two major subspecies of PE (36:5) detected in human plasma^23^. Soy-PC and Soy-PE displayed similar fatty acid composition, with linoleic acid (C18:2) and palmitic acid (C16:0) representing about 63-65.5 wt% and 14.9-17.7 wt% of total fatty acids, respectively (**Supplemental Figure 3A**), but differed in their head groups (choline vs ethanolamine) which allowed us to study the biological effects of the PE head group. By contrast, Soy-PE, ARA-PE and EPA-PE had the same head group but different fatty acid composition, which allowed us to study the biological effects of their fatty acid moieties (**Supplemental Figure 3A**).

We performed lipidomic analysis, to determine the precise phospholipid composition of the rHDLs. We confirmed that Soy-PC rHDL was composed mostly (99.98%) of PC alone, while Soy-PE rHDL was composed of PC and PE at the PC: PE molar ratio of 7.4:2.6. The phospholipidome of ARA-PE rHDL and EPA-PE rHDL was composed of PC with either ARA-PE at the PC: ARA-PE molar ratio of 7.2:2.8 or EPA-PE at the PC: EPA-PE molar ratio of 7.0:3.0, respectively (**Supplemental Figure 3B**). Such phospholipid composition is typical for rHDL prepared *in vitro*^33^ while the apoA-I: phospholipids molar ratio of 1:90 used (**Supplemental Table 2**) is close to that measured in human plasma HDL^14,42^.

### Presence of ARA-PE in rHDL diminished its ability to mitigate atherosclerosis in mice upon rHDL therapy

Although PE (36:5) exists in human plasma mainly as ARA-PE and EPA-PE, previous studies revealed a higher prevalence of ARA-PE in obese women with metabolic dysfunction and in patients with ASCVD^24,25^, suggesting that ARA-PE predominates in HDL from these patients. In order to test the impact of ARA-PE rHDL on atherosclerosis in mice, female Tg*CETP* x *Ldlr*^-/-^mice fed a high cholesterol diet were injected with either control (Soy-PC) or ARA-PE rHDL (**Figure 2A**). Analysis of the atherosclerotic plaques in the aortic root (**Figure 2B**) revealed a larger atherosclerotic lesion in mice injected with ARA-PE rHDL than in those injected with control Soy-PC rHDL (51.2% vs 41.0% of total aortic area, p=0.0005, respectively). This data indicates that the presence of ARA-PE in rHDL weakened the capacity of rHDLs to reduce atherosclerosis in mice and rendered them inefficient (compared with Supplemental figure 2D).

**Figure 2.**
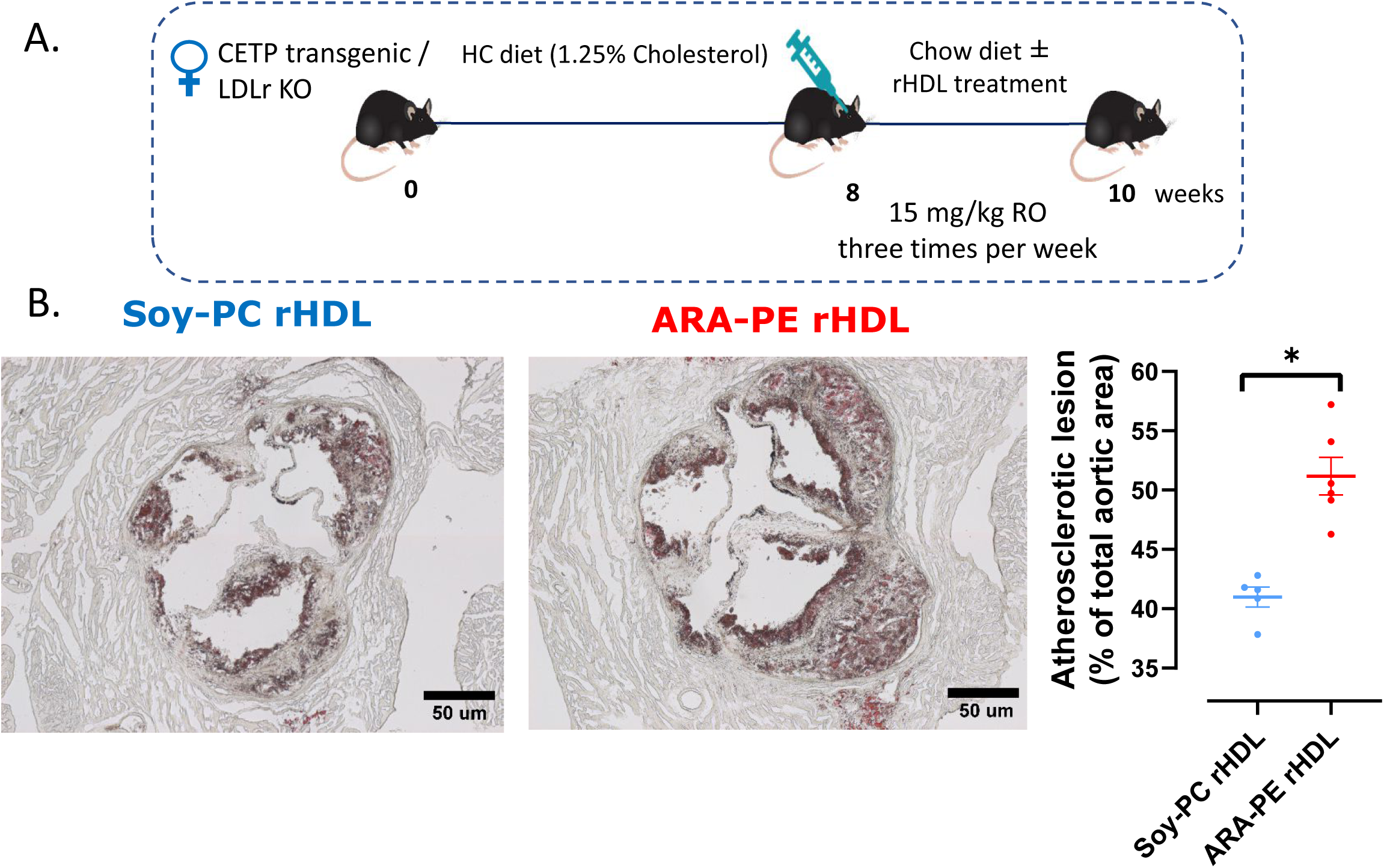
Injection of ARA-PE rHDL did not reduce the size of aortic atherosclerotic plaques as compared to Soy-PC rHDL in CETP transgenic-LDLr KO female mice. **A.** CETP transgenic/ LDL receptor KO female mice fed a high cholesterol diet for 8 weeks were injected with 15mg/kg of Soy-PC rHDL (n=5) or ARA-PE rHDL (n=6), retro-orbitally, every second day. The mice received 10 injections of either treatment, and were maintained on a chow diet during rHDL treatment. **B.** Sections of aortic root at the level of the three valves were labeled with hematoxylin-eosin to quantify the atherosclerotic plaque size. The extent of the atherosclerotic plaque was quantified using ImageJ software. Values are shown as mean (± SEM), n≥5, *p<0.001 vs. Soy-PC rHDL. CETP: cholesteryl ester transfer protein; HC: high cholesterol; rHDL: reconstituted high density lipoprotein; PC: phosphatidylcholine; ARE-PE: arachidonic acid-phosphatidylethanolamine.

### ARA-PE impaired the capacity of HDL to promote macrophagic cholesterol efflux while EPA-PE enhanced it

To determine whether the effects observed in humans and mice were accompanied by alterations in biological activities of HDL by to ARA-PE, we performed *in vitro* studies to evaluate the impact of PE and PE (36:5) in both ARA-PE and EPA-PE forms on the major cardioprotective functions of HDL. EPA-PE was included as the second major form of PE (36:5)^23^ to evaluate the reported potential beneficial effects of EPA on HDL functionality and atherosclerosis^29,30^. First, we explored the impact of the composition of rHDL particles (**Figure 3A**) on their cholesterol efflux capacity (CEC) from macrophages, which is the function of HDL proposed to underlie the association between low HDL-C levels and ASCVD^4,43^. As shown in **Figure 3B**, control Soy-PC rHDL exhibited a dose-dependent increase in CEC from human THP-1 macrophages, confirming the robust biological activity of this rHDL formulation. The CEC of Soy-PE rHDL was similar to that of Soy-PC rHDL and was only reduced at the highest concentration of rHDL at 50 µg/mL (−16%, p=0.015). Conversely, we observed a marked reduction of the CEC of ARA-PE rHDL in comparison to the both control Soy-PC and Soy-PE rHDLs at the concentrations of 20 µg/mL (−15% and −32%, respectively, p<0.05) and 50 µg/mL (−19% and −19%, respectively, p<0.05) and reduced by −15% (p<0.05) compared to Soy-PE rHDL at the concentration of 10 µg/mL. Interestingly, such alterations of the CEC were not detected in EPA-PE rHDL; this metric was rather greatly enhanced by an average of +87% (p<0.05) as compared to the other rHDL particles. The quantification of mRNA levels by qPCR as well as of the ABCA1 surface expression by flow cytometry in human THP-1 macrophages following incubations with the rHDLs (**Supplemental Figure 4A-C**) suggested that the opposite effects of ARA-PE and EPA-PE on the CEC of rHDL were not due to a modification of the expression of the major lipid transporters involved in cholesterol efflux. Taken together, these findings indicate that the presence of ARA-PE impaired the CEC of rHDL in human macrophages while that of EPA-PE improved it.

**Figure 3.**
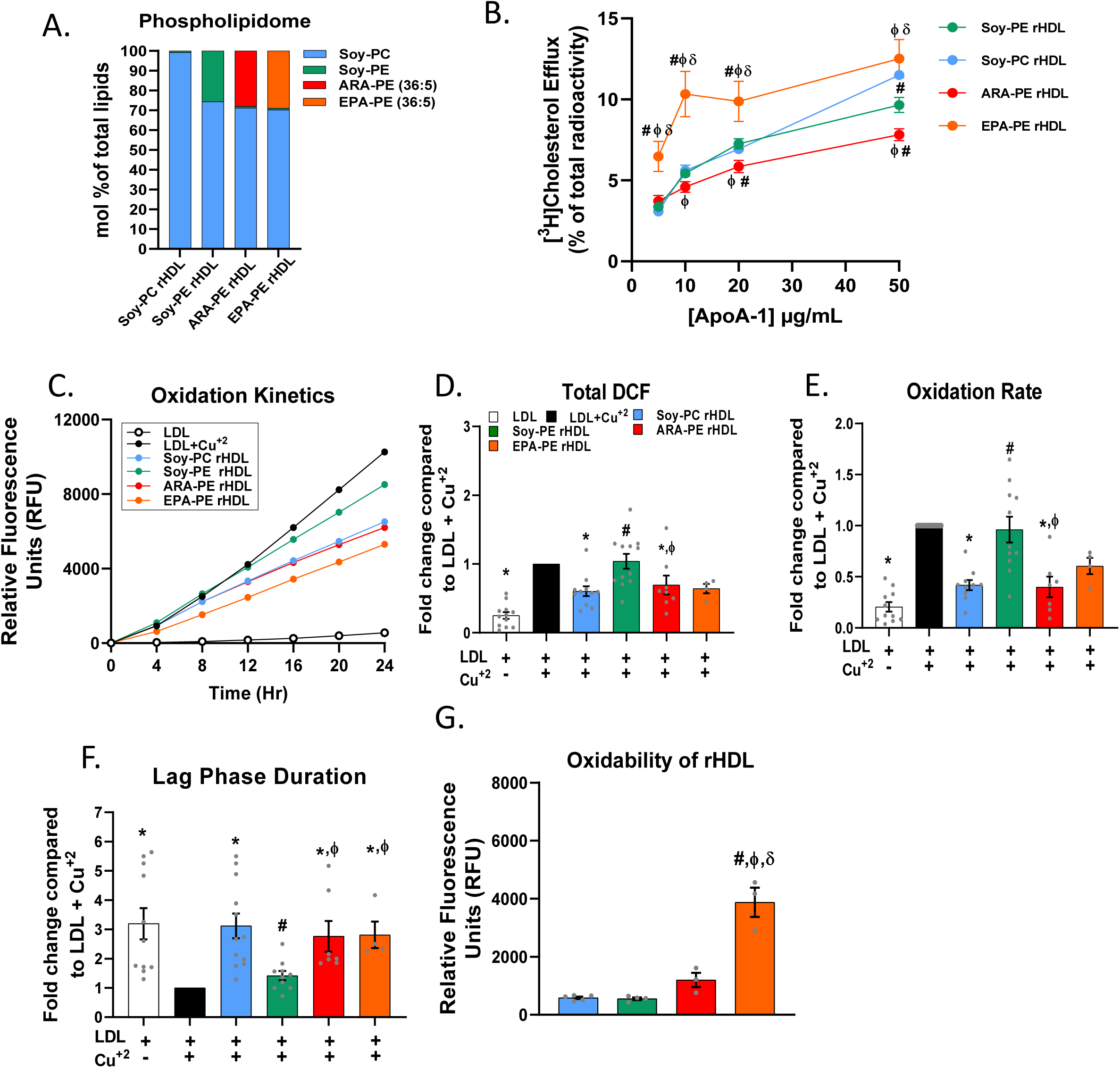
The presence of Soy-PE, ARA-PE or EPA-PE alters the biological activities of Soy-PC rHDL. **A.** The phospholipidome composition of the rHDLs determined by LC/MS-MS. **B**. The CEC of rHDLs was determined using THP-1 macrophages loaded with radiolabeled [^3^H] cholesterol and incubated with 5, 10, 20, or 50 µg ApoA-1/ml of Soy-PC, Soy-PE, ARA-PE, and EPA-PE rHDLs for 4 hours. **C.** DCFH fluorescent probe was used to assess the oxidation kinetics of LDL induced by copper sulfate (CuSO_4_) in the presence of Soy-PC, Soy-PE, ARA-PE, or EPA-PE rHDLs (50µg ApoA-I/ml), or none for 24 hours at 37C° using Spectromax Gemini fluorometer. **D.** Maximum DCF (oxidized DCFH) formed after 24 hours. **E.** Oxidation rate during the propagation phase. **F.** Lag phase duration, **G.** Oxidability of rHDL expressed as relative fluorescence units (RFU) for different rHDLs (50µg ApoA-I/ml), incubated with CuSO4, and DCFH (0.2mg/ml) in the absence of LDL. *vs. LDL+Cu^+2^, ^#^vs. Soy-PC rHDL, ^φ^vs. Soy-PE rHDL ^δ^vs. ARA-PE rHDL with p values <0.05. Values are shown as mean ±SEM from at least 3 independent experiments that were performed in triplicate for each condition.

### EPA and ARA counteract the abolishment of the AOX of rHDL mediated by PE

We next investigated the capacity of the PE-containing rHDLs to prevent LDL oxidation, which is a key element of the pathogenesis of atherosclerosis^2^. The antioxidative activity (AOX) of rHDLs was evaluated as their capacity to prevent copper-induced LDL oxidation using a DCFH fluorescent probe. The extent of LDL oxidation was evaluated using a biphasic oxidation kinetics (**Supplemental Figure 5**), including the maximum oxidized DCFH formed at the end of the experiment and the oxidation rate in the propagation phase which are proportional to the level of LDL oxidation, as well as the duration of the lag phase which is inversely related to the level of LDL oxidation. In accordance with oxidation kinetics in **Figure 3C**, we observed that control Soy-PC rHDL efficiently reduced both the maximum oxidized DCFH (−33%, p=0.004, **Figure 3D**), and the oxidation rate in the propagation phase (−58%, p<0.0001, **Figure 3E**), as well as prolonged the lag phase 2.7-fold (p<0.0001, **Figure 3F**) compared to LDL only treated with copper, indicating an efficient AOX of control Soy-PC rHDL. Strikingly, this AOX was lacking with Soy-PE rHDL but conserved when either ARA-PE or EPA-PE rHDLs were used (**Figure 3D-F**), suggesting that ARA and EPA may restore the AOX abolished by Soy-PE rHDL.

Several studies reported that HDL may protect LDL from oxidation in part by acting as a sacrificial target for oxidation^44^. Thus, we investigated the oxidability of rHDLs by copper ions in the absence of LDL. Our results indicated was EPA-PE rHDL was significantly more susceptible to oxidation than all other rHDLs (**Figure 3G**). Overall, our findings demonstrate that Soy-PE abolished the AOX of rHDL but this effect could be reversed by EPA and ARA.

### EPA and ARA restored the PE-mediated alteration of the phospholipid transfer from LDL to rHDL by increasing its lipid fluidity

Because the first step in the AOX of HDL is thought to involve the transfer of oxidized phospholipids from LDL to HDL in a process controlled by the fluidity of lipids at the surface of HDL^45^, we investigated the capacity of rHDLs to acquire phospholipids from LDL labelled with a Dil fluorescent probe. As shown in **Figure 4A**, the transfer of the fluorescent phospholipid from DiI-labelled LDL to control Soy-PC rHDL was similar to that to reference normolipidemic HDL (employed as ApoB-depleted plasma), highlighting the efficient capacity of rHDL to accept phospholipids. However, the transfer of fluorescent phospholipid from DiI-labelled LDL to Soy-PE rHDL was significantly reduced (−26%, p=0.0003). By contrast, the transfer was increased to EPA-PE and ARA-PE rHDLs when compared to both control Soy-PC rHDL and Soy-PE rHDL (EPA-PE: +42%, p=0.005 and +91%, p=0.0006; ARA-PE: +27%, p=0.026 and +72%, p=0.001, respectively). In an attempt to gain mechanistic insights into the impact of PE and PE (36:5) on HDL fluidity, we investigated the surface and core lipid fluidity of rHDL using TMA-DPH and DPH fluorescent probes^39^, which are located in the surface and core region of HDL, respectively. Fluorescence anisotropy values measured for Soy-PE rHDL and ARA-PE rHDL with TMA-DPH (**Figure 4B**) and DPH (**Figure 4C**) probes were not statistically different from those obtained for control Soy-PC rHDL. By contrast, the anisotropy values of the TMA-DPH and DPH probes were both decreased in EPA-PE rHDL (−46%, p=0.02, −96%, p=0.0017, respectively) compared to control Soy-PC rHDL (and to ARA-PE rHDL for the DPH probe), collectively indicating an increased surface and core luidity of EPA-PE rHDL.

**Figure 4.**
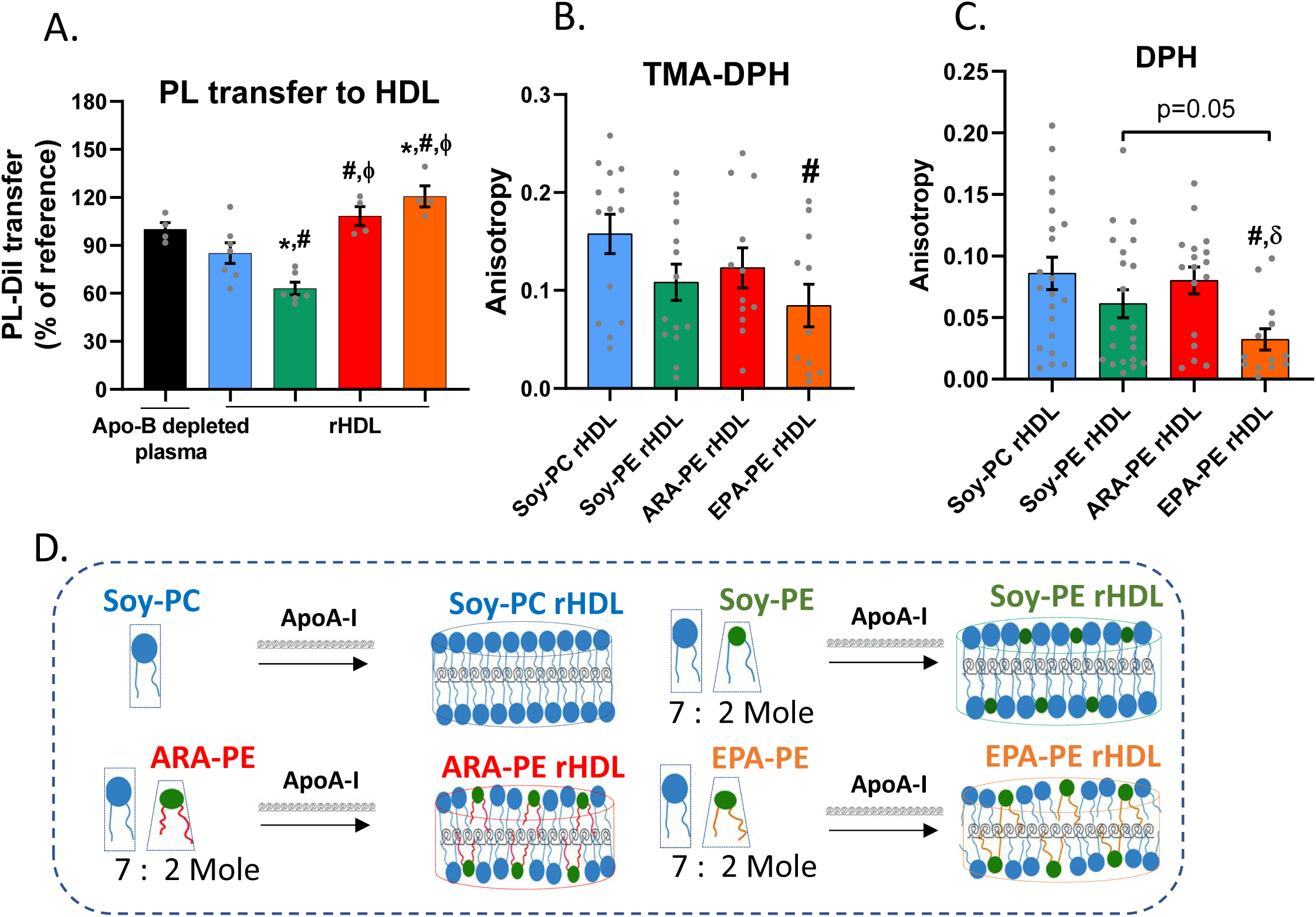
Soy-PE diminished the capacity of rHDL to accept phospholipids from LDL, while ARA-PE and EPA-PE reversed this effect, potentially by enhancing the fluidity of HDL lipids. **A.** The percentage of Dil PL transferred from DiI labelled LDL to rHDLs. To evaluate the capacity of rHDLs to acquire phospholipids (PL), Dil-Labelled LDL was mixed with rHDL (4 mg ApoA-I/dL), or reference ApoB deficient plasma (1:30 v/v), and incubated for 1 hour at 37°C to achieve PL transfer. Then, mixtures were ApoB-depleted and the fluorescence intensity of Dil was measured in rHDL. For the fluidity assay, DPH or TMA-DPH were incorporated in rHDL particles, and the anisotropy of the probe fluorescence was recorded. **B.** Fluorescence anisotropy of TMA-DPH probe, which is inversely related to surface lipid fluidity of rHDL. **C.** Fluorescence anisotropy of DPH probe, which is inversely related to core lipid fluidity of rHDL. **D.** Scheme illustrates the influence of Soy-PE, ARA-PE and EPA-PE on the arrangement of the phospholipid monolayer of rHDLs, according to their head group and the unsaturation level in their fatty acid moieties. Our data indicate that Soy-PE reduced the capacity of rHDL to accept phospholipids from LDL, while the fatty acid moieties of ARA-PE and EPA-PE enhanced it, which could result from their fluidizing effects. In addition, these data point out that EPA-PE rHDL represented the most fluid particle of all the rHDLs studied. *vs. ADP, ^#^vs. Soy-PC rHDL, ^φ^vs. Soy-PE rHDL, ^δ^vs. ARA-PE rHDL, with p values <0.05. Values are shown as mean±SEM from at least 3 independent experiments that were performed in triplicate for each condition. TMA-DPH: trimethylamine-diphenylhexatriene; DPH: diphenylhexatriene.

Taken together, these findings led us to propose that the loss of the AOX in PE-containing rHDL could result from an alteration of the phospholipid transfer from LDL. On the contrary, such a transfer of oxidized phospholipids was enhanced when PE were enriched in EPA and ARA. It was noteworthy that EPA exhibited the strongest fluidizing effect by enhancing both surface and core fluidity of rHDL (**Figure 4D**) suggesting a mechanism for the enhanced transfer of oxidized phospholipids to EPA-PE rHDL involving a fluidization of rHDL lipids.

### EPA but not ARA rescued the anti-inflammatory activity of PE-containing rHDL in macrophages upon LPS treatment

Because HDL exert potent anti-inflammatory activities (AIA) allowing a reduction of inflammatory activation of arterial macrophages^2^, we investigated the influence of a short-term (ST, 4h) and long-term (LT, 16h) treatment with rHDLs on cellular production of pro-inflammatory IL-1β cytokine whose inhibition was reported to reduce cardiovascular events in patients with CVD^46^. In human THP-1 macrophages, mRNA levels of IL-1β triggered by LPS were reduced following both ST (−37%, p=0.03; **Figure 5A**) and LT (−30%, p=0.001; **Figure 5B**) pre-incubation with control Soy-PC rHDL. The inhibitory effect of the ST and LT pre-incubation with rHDL was no longer observed with either Soy-PE or ARA-PE rHDLs. Noteworthy, secretion was significantly reduced following the ST (−63%, p=0.01; **Figure 5A**) and LT (−50%, p=0.04; **Figure 5B**) pre-incubation with EPA-PE rHDL. It is of note that rHDLs were without effect on the cell surface expression of the LPS binding receptor TLR4 (**Supplemental Figure 4E**).

**Figure 5.**
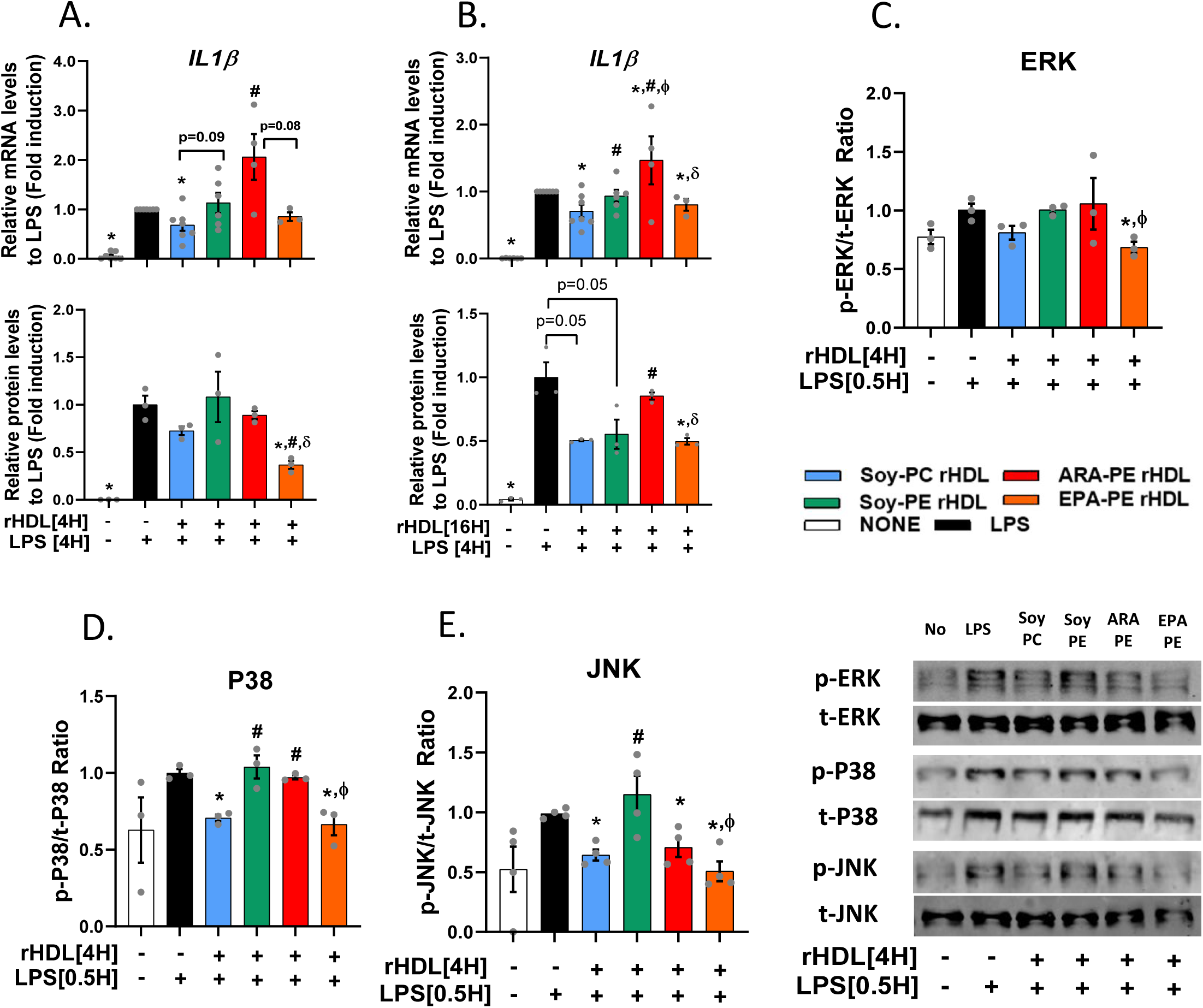
Soy-PE reduced the AIA of rHDL, while ARA-PE endowed rHDL with pro-inflammatory activity with EPA-PE reversing it and improving the AIA of rHDL. AIA of rHDLs was evaluated in LPS-stimulated THP-1 macrophages **A.** Relative mRNA and protein levels of IL-1β after short-term (4 hours), and **B.** long-term (16 hours) treatment of THP-1 cells with 20 µg ApoA-I/mL of different rHDL followed by 4 hours stimulation with 100 ng/mL of LPS. **C-E.** Western blot analysis for the phosphorylated and non-phosphorylated ERK, p38, and JNK MAPKs in THP-1 cells treated with rHDLs (20 µg ApoA-I/mL) of for 4 hours, followed by 0.5hour stimulation with 100 ng/mL of LPS. *vs. LPS, ^#^vs. Soy-PC rHDL, ^φ^vs. Soy-PE rHDL, ^δ^vs. ARA-PE rHDL, with p values <0.05. Values shown are from at least 3 independent experiments that were performed in triplicate for each condition. IL-1β: Interleukin-1 beta; LPS: lipopolysaccharide; JNK: Jun kinase; ERK: Extracellular signal-regulated kinase; MAPK: mitogen-activated protein kinase.

To test if the anti-inflammatory effect of rHDLs involved modulation of intracellular inflammatory signaling pathways in macrophages, we investigated their impact on the LPS-induced activation of ERK, p38 MAPK and JNK signaling pathways which were previously reported to be downregulated in macrophages by HDL^47^. Control Soy-PC rHDL reduced the LPS-induced phosphorylation of p38 MAPK and JNK in human THP-1 macrophages (−30%, p=0.001 and −36%, p=0.002, respectively, **Figure 5C-E**). Such inhibitory effects were abolished with Soy-PE rHDL and ARA-PE rHDL, with the exception of JNK phosphorylation which was reduced with ARA-PE rHDL to the same extent as with control Soy-PC rHDL. By contrast, the reduction of the LPS-induced phosphorylation of p38 and JNK was retained by EPA-PE rHDL (−32%, p=0.01; −34%, p=0.01 and −50%, p=0.008, respectively). Moreover, the presence of EPA-PE in rHDL made them able to decrease the phosphorylation of ERK.

These findings demonstrate that the presence of PE and ARA-PE abrogated the anti-inflammatory activity of rHDL in LPS-treated human macrophages by acting on inflammatory signaling pathways and that the presence of EPA was able to restore it.

### Composition of PE-containing rHDL in ARA and EPA played an important role in eicosanoid production and IL-1 β secretion in human macrophages

We finally addressed the hypothesis that ARA carried by HDL could promote inflammation in macrophages by providing a source of ARA which could be oxidized to produce pro-inflammatory eicosanoids. As shown in **Figure 6A**, 16h-treatment of human THP-1 macrophages with ARA-PE rHDL was accompanied by a marked increase of the production of eicosanoids, including hydroxyeicosatetraenoic acids (5S-, 11S-, and 15S-HETEs), prostaglandin (PGE_2_) and thromboxane (TXB_2_), in comparison to control Soy-PC and Soy-PE rHDLs. Substitution of ARA by EPA in PE-rHDL partly abolished this effect and led to a marked decrease in 5S-HETE compared to control Soy-PC rHDL and both Soy-PE rHDL and ARA-PE rHDL. Interestingly, the impact of PUFAs (ARA and EPA) in PE-rHDLs on eicosanoids production was associated with a concomitant effect on the secretion of pro-inflammatory IL-1β cytokine by human THP-1 macrophages (**Figure 6B**). Strikingly, whereas treatment with ARA-PE rHDL led to a 5-fold increase of IL-1β secretion in comparison to control Soy-PC and Soy-PE rHDLs, the secretion of IL-1β was almost undetectable in macrophages incubated with EPA-PE rHDL.

**Figure 6.**
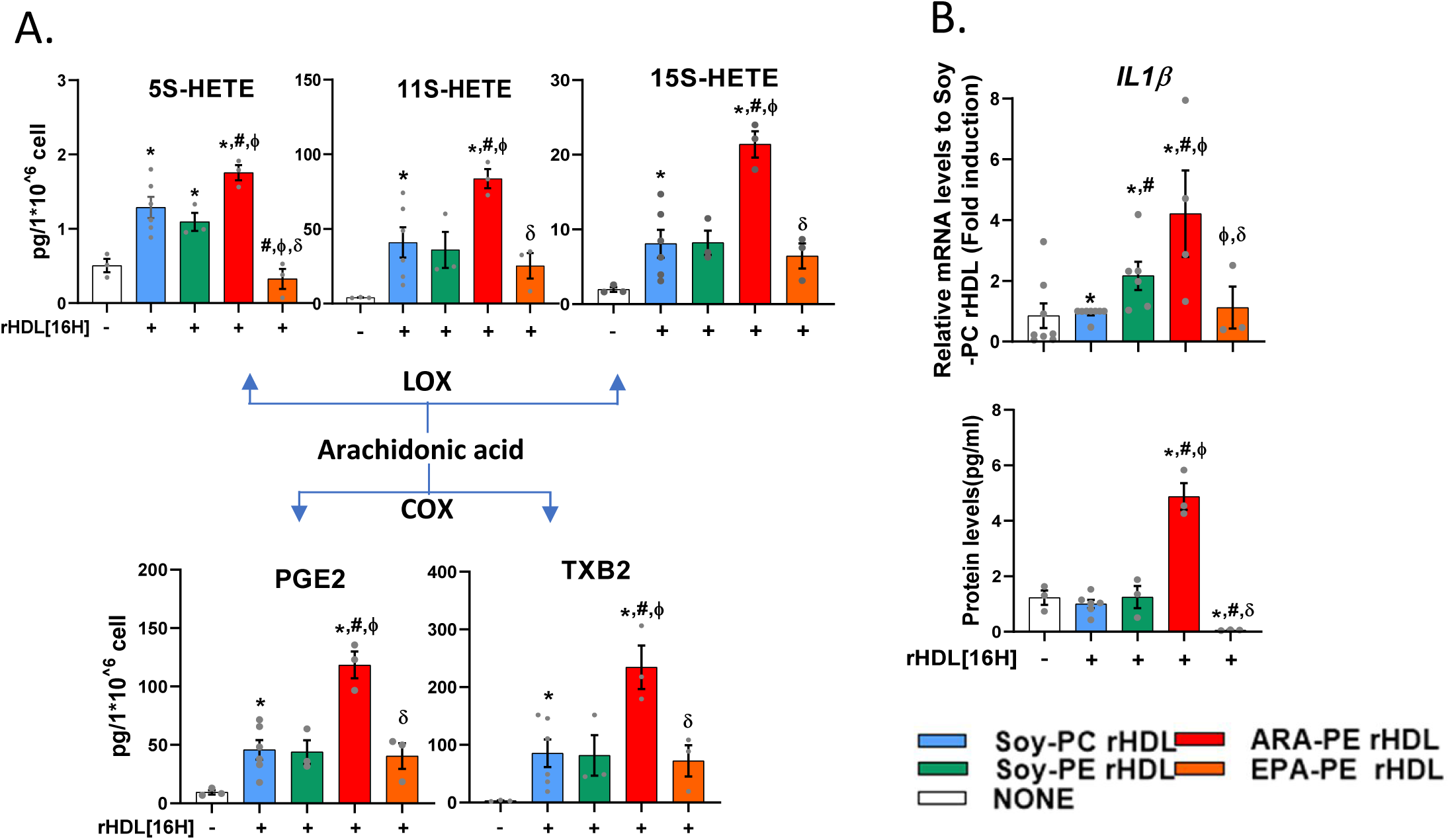
ARA-PE exacerbated Soy-PE rHDL pro-inflammatory effects while EPA-PE reversed it and even improved the basal AIA of rHDL. **A.** Pro-inflammatory eicosanoid production after 16 hours treatment of THP-1 cells with 20µg/mL of rHDLs (20µg APOA-1/mL). The pro-inflammatory effects of rHDLs were evaluated in THP-1 macrophages in the absence of LPS stimulation. **B.** Relative mRNA and protein levels of IL-1β after long-term (16 hour) treatment of THP-1 cells with 20 µg ApoA-I/mL of rHDL.*vs. None, ^#^vs. Soy-PC rHDL, ^φ^vs. Soy-PE rHDL, ^δ^vs. ARA-PE rHDL, with p values less than 0.05. Values are from at least 3 independent experiments that were performed in triplicate for each condition. HETE: hydroxyeicosatetraenoic acids; PGE2: prostaglandin E2, TXB2: thromboxane B2, LOX: lipoxygenase, and COX: cyclooxygenase enzymes.

These results provide evidences that the ARA in PE-containing rHDLs contributed to the production of eicosanoids and the secretion of the pro-inflammatory IL-1β cytokine by human macrophages. By contrast, the EPA content in PE-containing rHDL exhibited a potent inhibitory action toward IL-1 β secretion from human macrophages.

## Discussion

Plasma lipidomic studies showed a robust positive association between several PE species, notably PE (36:5), with ASCVD^16,18^ although the underlying mechanisms are unknown. We demonstrated that the contents of total PE and PE (36:5) in large HDL are linked to atherosclerosis development in MetS. The atherogenic effect of ARA-PE was confirmed in mice injected with rHDL containing ARA-PE, the predominant form of PE in patients with cardiometabolic diseases^24,25^. Our *in vitro* investigations revealed that the presence of PE in rHDL impaired their biological activities, an effect worsened with ARA-PE, suggesting deleterious roles of the presence of the both PE head group and ARA lipid chain for anti-atherogenic functions of HDL with potential implications for ASCVD pathogenesis. On the contrary, EPA-PE counteracted the PE-mediated rHDL dysfunction and enhanced the anti-atherogenic functions of rHDL, adding a novel element to mechanistic insights in the protective role of EPA in patients with ASCVD,

In agreement with our findings in MetS patients, Giraud et al. reported increased levels of ARA-containing PE species in HDL from patients with rheumatoid arthritis; their abundance was associated with inflammation and CVD risk^48^. Taken together with the present study, these data suggest that the association of plasma PE (36:5) with ASCVD may result in part from the presence of ARA-PE, i.e. PE (16:1/20:4), in HDL, rather than EPA-PE, i.e. PE (16:0/20:5), the other form of PE (36:5) detected in human plasma^23^. This hypothesis is reinforced by the numerous studies reporting the positive link of ARA (C20:4), a key omega-6 PUFA, with ASCVD^24^, which contrasts multiple anti-atherogenic properties of EPA (C20:5), an omega-3 PUFA^49^. Although our findings undoubtfully warrant further validation in larger cohorts of patients with ASCVD, the atherogenic property of ARA-PE rHDL was validated in our *in vivo* experiments in a mouse model of atherosclerosis injected with rHDL. Indeed, we found that ARA-PE rHDL lacked a capacity to reduce atherosclerosis compared to control rHDL, underlying the deleterious role of ARA-PE on HDL-mediated atheroprotection.

The present study demonstrated that PE and ARA-PE impaired several biological activities of rHDL particles proposed to contribute to their atheroprotective action. Thus, the presence of Soy-PE (a mixture of PE species) led to an attenuation of the CEC of rHDL from human macrophages in comparison to Soy-PC rHDL with a similar fatty acid composition. This results is coherent with a previous study showing an inverse correlation between the total PE content of HDL and the capacity of human serum to promote cellular cholesterol efflux^22^. Because mRNA levels of transporters and receptors controlling cholesterol efflux from macrophages, such as ABCA1, the major cholesterol transporter in human macrophages^36,42^, were not altered by PE-containing rHDL, these data support an extracellular rather than an intracellular influence of PE on the CEC of HDL. This hypothesis is consistent with the work of Demel et al. who described a lower affinity of PE compared to PC toward cholesterol^50^, suggesting a reduced membrane cholesterol transfer into PE rHDL. Moreover, PE can alter the dynamic structure and interaction of HDL with cell membrane receptors like ABCA1^21,51^. Taken together, these mechanisms may contribute to the capacity of PE to impair CEC of rHDL. Interestingly, a more pronounced impairment in the CEC of PE-containing rHDL was observed in the presence of ARA-PE, suggesting additional deleterious effects mediated by ARA. Although the mechanism underlying such a role for ARA was not identified in our study, it is noteworthy that the enrichment of HDL with another omega-6 fatty acid, i.e. linoleic acid, similarly led to a reduced CEC of HDL^31^.

In addition to the effects on CEC, our study demonstrated that the addition of Soy-PE diminished the AOX of rHDL. Seeking for mechanistic insights, we observed that PE-containing rHDL exhibited a reduced capacity to acquire phospholipids from LDL compared to control rHDL which could account for its diminished AOX. Indeed, antioxidative function of HDL toward LDL requires the transfer of oxidized phospholipids, including lipid hydroperoxides, from LDL to HDL, which is enhanced by HDL’s surface fluidity, followed by their reduction via redox-active methionine (Met) residues of apoA-I^45,52^. PC- and PE-containing rHDL used in our *in vitro* experiments only differed in their head groups (choline vs ethanolamine). Ethanolamine head groups are considered to be more compact than choline^53^, due to their capacity to form non-covalent bonds with neighboring lipids thereby restricting their movement^53,54^. This rigidity could limit the incorporation of exogenous phospholipids into PE rHDL by increasing interactions with adjacent lipids on the HDL surface as observed in our experiments. Several reports indicated that PE has limited interactions with ApoA-I and can alter both the content and conformation of ApoA-I in rHDL^21,51^. This could contribute to the diminished AOX of our PE rHDL by affecting the availability of apoA-I Met residues and HDL interactions with other lipoproteins, like LDL^21^. In contrast, replacing Soy-PE with ARA-PE preserved the AOX of rHDL, suggesting that ARA may restore the impaired AOX of PE-rHDL. In this regard, we observed an enhanced transfer of phospholipids from LDL to ARA-PE rHDL, likely reflecting its increased surface fluidity which could potentially account for its conserved AOX.

Finally, our data on the influence of PE and ARA-PE on the AIA of Soy-rHDL in pro-inflammatory human macrophages treated with LPS demonstrated that the presence of PE abolished the capacity of rHDL to decrease the expression and secretion of IL-1β. Previous studies have established a strong link between CEC and AIA of HDL^2,47^, suggesting that the impaired AIA of PE-containing rHDL could result from its impaired CEC. In this regard, Sun et al. linked cholesterol accumulation in plasma membrane of macrophages with enhanced TLR-4 signaling and p38 MAPK activation^55^. Although cell surface expression of TLR4 in human macrophages was not altered in the present study by the presence of PE in rHDL, the phosphorylation of p38 MAPK and activation of other inflammatory signaling pathways (ERK, JNK) known to be modulated by HDL^47^ was no longer attenuated by PE-containing rHDL, providing mechanistic clues on the negative impact of PE on the AIA of HDL.

In human macrophages non-treated with LPS, the presence of PE converted rHDL into pro-inflammatory particles, which are recognized as contributors to inflammation in atherosclerosis^56^. Interestingly, although the enrichment of PE with ARA has limited additional effect on the pro-inflammatory effects of PE-rHDL in LPS-treated human macrophages, a potent increase in the expression and secretion of IL-1β was detected in non-treated cells following the long-term incubation with ARA-PE rHDL. We propose that the proinflammatory effect of ARA-PE rHDL mainly resulted from an increased production of eicosanoids, including HETEs (5S, 11S, and 15S), PGE2, and TXB2, which were found elevated in the present study when macrophages were incubated with ARA-PE rHDL in comparison to others rHDL particles. Indeed, ARA is the primary precursor for the pro-inflammatory eicosanoids which are potent lipid mediators implicated in the inflammatory process of ASCVD^24,28^. Eicosanoid generation mainly involves ARA release from cell membrane phospholipids by phospholipase A2 followed by enzymatic metabolism of ARA by cyclooxygenases (COXs) generating PGs and TXs, and by lipoxygenases (LOXs) generating HETEs and leukotrienes^28^. Our findings uncovered that HDL, primarily PE-containing HDL, can be a source of ARA in macrophages for inducing inflammatory response through eicosanoid production. Such a mechanism could play a role in cellular IL-1 β secretion and the development of ASCVD.

Although omega-3 PUFAs, including the mixtures of EPA and DHA, were without effects on cardiovascular events in large clinical trials ^57–59^, a highly purified EPA ethyl ester was reported to reduce the CV risk^26^, suggesting a specific cardioprotective effect of EPA that would not be exerted by other omega-3 PUFAs. Our findings thereby provide new mechanistic insights regarding the potential impact of EPA on ASCVD. Indeed, our data demonstrated that EPA was able to counteract the deleterious effects of PE, especially those of ARA-PE, on rHDL functions. Thus, EPA-PE restored the altered AOX of PE-containing rHDL to a higher degree than did ARA-PE. The positive influence of omega-3 PUFA, including EPA, on the AOX of both rHDL and plasma HDL has been well-documented^30,31^. This effect can be attributed to the EPA structure composed of an additional double bond compared to ARA, rendering rHDL more fluid and improving their capacity to acquire and inactivate oxidized lipids. Moreover, we observed that EPA-PE enhanced both surface and core fluidity of rHDL, this latter parameter being only increased by EPA-PE rHDL under our experimental conditions.

Our study also demonstrated that EPA-PE restored the anti-inflammatory activity of rHDL in LPS-treated macrophages which was abolished in PE-containing rHDL. Strikingly, a similar effect of EPA-PE rHDL on the inhibition of IL-1β secretion was also observed in the absence of LPS when control rHDL was without effect on inflammatory processes in macrophages. The well-established anti-inflammatory effects of EPA^24,49,60^, especially its capacity to attenuate LPS-induced IL-1β secretion by macrophages^61^, could contribute to the strong AIA of EPA-PE rHDL. Tian et al. reported that dietary EPA-PC and EPA-PE attenuated the inflammatory phenotype of macrophages by reducing IL-1β expression and stimulating M2 anti-inflammatory macrophage polarization^62^ through a role of EPA in eicosanoid production^63^. In agreement with this report, we found that EPA-PE rHDL not only abolished the increased production of eicosanoids observed with ARA-PE rHDL, but also markedly reduced 5S-HETE levels in human macrophages compared to both Soy-PC- and Soy-PE-containing rHDL, further supporting the importance of PUFA composition in PE HDL for the production of eicosanoids in macrophages.

Finally, the presence of EPA-PE improved the cholesterol efflux capacity of rHDL from macrophages, a metric of HDL function inversely associated with ASCVD^4–6^. A similar observation was reported by Tanaka et al. with EPA-PC enhancing the CEC of rHDL compared to PC rHDL^29^, suggesting that the effect of EPA on the CEC of HDL can be independent of the nature of PL classes. In this context, the elevation of both the surface and core lipid fluidity of rHDL by EPA-PE compared to PC-containing rHDL could potentially account for the enhanced CEC by facilitating cholesterol insertion at the particle surface. This hypothesis is consistent with the established role of rHDL surface fluidity as a key determinant of this process^10,64^. Moreover, the increased core fluidity of EPA-PE rHDL could further facilitate cholesterol incorporation into rHDL. Such profound fluidizing effects of EPA-PE are likely due to the high unsaturation level of EPA, conferring a better structural flexibility for adopting highly kinked conformations in the phospholipid monolayer of rHDL^65^.

In conclusion, our study provides evidence that PE species, particularly ARA-PE, contribute to the formation of dysfunctional HDL, consistent with the deleterious role of PE (36:5) in ASCVD reported by epidemiological studies. Circulating levels of PE (36:5) species, i.e. ARA-PE, might therefore serve as novel biomarkers of HDL functionality while our observations on EPA-containing rHDL can provide new approaches to the formulation of reconstituted HDL for therapeutic use in ASCVD^3^. Importantly, our data provide mechanistic insights on the cardioprotective role of EPA in ASCVD by counteracting the PE-mediated dysfunction of HDL. Such a role might contribute to the reduction of cardiovascular events observed in patients consuming highly purified EPA ethyl ester, such as in the REDUCE IT clinical trial^26^, supporting the use of EPA-rich diets as a therapeutic strategy to enhance HDL-mediated cardiovascular protection.

## Abbreviations

Apo: apolipoprotein
ARA: arachidonic acid
ASCVD: atherosclerotic cardiovascular disease
CAD: coronary artery disease
CETP: cholesteryl cster transfer protein
cIMT: carotid intima-media thickness
CMD: cardiometabolic diseases
DPH: diphenylhexatriene
EDTA: ethylenediaminetetraacetic acid
EPA: eicosapentaenoic acid
ER: endoplasmic reticulum
FPLC: fast protein liquid chromatography
HDL-C: high-density lipoprotein
IL-1β: interleukin-1 beta
LA: linoleic acid
LPC: lysophosphatidylcholine
LPDP: lipoprotein-deficient plasma
LPS: lipopolysaccharide
Met: methionine
MetS: metabolic syndrome
PBS: phosphate buffer saline
PC: phosphatidylcholine
PE: phosphatidylethanolamine
PI: phosphatidylinositol
PLOOH: phospholipid hydroperoxide
PUFA: polyunsaturated fatty acid
rHDL: reconstituted HDL
Soy-PC: L-α phosphatidylcholine
Soy-PE: L-α phosphatidylethanolamine
TMA-DPH: trimethylamine-diphenylhexatriene

## Acknowledgements

The authors are indebted to all the participants for their cooperation.

## Sources of funding

This study was supported by French National Institute for Health and Medical Research (INSERM) and Sorbonne Université (Paris, France). D.M. and W.L.G. acknowledge support from the Agence Nationale pour la Recherche (ANR-19-CE14-0020). W.L.G. acknowledges supports from the Fondation de France (00066330), Alliance Sorbonne Université (Programme Emergence) and the Société Francophone du Diabète. LMH was a recipient of a doctoral contract from CORDDIM. VDD and CR were recipients of doctoral contract from Sorbonne Université. CM was a recipient of a doctoral contract from the Fondation pour la Recherche Médicale (FRM). MT was a recipient of a doctoral contract from Arab American University of Palestine (Jenin, Palestine).

## Disclosures

The authors declare to have nothing to disclose.

## Highlights

- This study identifies arachidonic acid-phosphatidylethanolamine (ARA-PE) as a key contributor to HDL dysfunction, which is closely linked to the development of atherosclerosis.
- Replacement of ARA by eicosapentaenoic acid (EPA) can counteract this effect by restoring HDL’s protective functions.
- EPA-rich diets or reconstituted HDL containing EPA-PE, can serve as innovative approaches for reducing atherosclerotic cardiovascular disease.

## Notes

### Competing Interest Statement

The authors have declared no competing interest.

